# Long non-coding RNA small nucleolar RNA host genes as prognostic molecular biomarkers in hepatocellular carcinoma: a meta-analysis

**DOI:** 10.1101/2023.05.07.539768

**Authors:** Meng Huang, Zhiwen Zhao, Lihua Yang

## Abstract

**Objective:** Recently, increasing data have suggested that the lncRNA small nucleolar RNA host genes (SNHGs) were aberrantly expressed in hepatocellular carcinoma(HCC), but the association between the prognosis of HCC and their expression remained unclear. The purpose of this meta-analysis was to determine the prognostic significance of lncRNA SNHGs in HCC.

**Methods:** We systematically searched Embase, Web of Science, PubMed, and Cochrane Library for eligible articles published up to October 2022. The prognostic significance of SNHGs in HCC was evaluated by hazard ratios (HRs) and 95% confidence intervals (CIs). Odds ratios (ORs) were used to assess the clinicopathological features of SNHGs.

**Results:** This analysis comprised a total of 25 studies covering 2314 patients with HCC. The findings demonstrated that over-expressed SNHGs were associated with larger tumor size, multiple tumor numbers, poor histologic grade, earlier lymphatic metastasis, vein invasion, advanced tumor stage, portal vein tumor thrombosis (PVTT), and higher AFP level, but not with gender, age, HBV infection, and cirrhosis. In terms of prognosis, patients with higher SNHG expression were more likely to have shorter overall survival (OS), relapse-free survival (RFS), and disease-free survival (DFS).

**Conclusions:** In conclusion, upregulation of SNHG expression correlated with clinicopathological parameters and could predict a poor prognosis for HCC patients.

## 1. Introduction

According to 2020 Cancer Data, hepatocellular carcinoma (HCC) is the sixth most prevalent cancer and the third leading cause of cancer death worldwide, resulting in a significant disease burden [1]. Metabolic risk factors for HCC are becoming more prevalent despite the hepatitis virus remaining the main cause [2]. At the moment, non-drug treatments include ablation, transarterial chemoembolization (TACE), liver transplantation, and hepatic resection. In the meanwhile, advanced HCC is primarily treated systemically with medications including monoclonal antibodies like nivolumab and small molecule targeted medications like sorafenib and lenvatinib [3]. Most patients are diagnosed in the middle or late stages, with a 5-year survival rate below 18% [4]. However, current therapy advances still fail to provide satisfactory outcomes. Therefore, it is critical and vital to develop effective biomarkers early on that can be applied as targeted therapy for HCC.

LncRNAs are non-coding RNAs with a length greater than 200 nucleotides. In the absence of functional open reading frames, they hardly encode any protein [5]. Increasing studies have clarified that lncRNAs, initially considered as genomic junk, can act as oncogenes or antioncogenes in cancer, regulating tumor growth, metastasis, metabolism, and progression, and may offer a new method for cancer diagnosis and treatment [6]. Multiple lncRNAs about HCC have been found to exhibit abnormal expression and take a role in malignant phenotypes via binding to DNA, RNA, or proteins or by encoding tiny peptides, which could influence the progression of HCC. They were expected to be potential biomarkers for diagnosis and prognosis of HCC [7]. Long non-coding small nucleolar RNA host genes (lnc-SNHGs) are host genes for snoRNAs (small nucleolar RNAs), which are overexpressed in human cancers. When acting as a lncRNA called SNHG, snoRNAs retain full-length transcripts, including exons [8]. Five different molecular mechanisms explain how SNHGs work: three are connected to cytoplasmic localization, reducing miRNA bioavailability through molecular sponge activity, limiting translation, and preventing ubiquitination; the other two are related to nuclear localization, interacting with transcription factors or repressors and altering DNA methylation. SNHGs are crucial in carcinogenesis and cancer progression through influencing DNA, RNA, and protein [9]. Additionally, SNHGs take part in the occurrence, growth, and pathophysiology of gastrointestinal cancers by regulating downstream targets. They also correlate with clinicopathological traits such as shorter overall survival, tumor size, lymph node metastasis, and TNM stage [10]. It has been shown that upregulating the expression of SNHGs suppressed tumor growth and was strongly associated with clinicopathological features. According to an examination of experimental data, HCC patients expressing high levels of SNHGs got a worse prognosis. Various studies have confirmed their predictive value, although it is unclear how each of them affects the prognosis of HCC [11]. In 2017, LI et al. [12] published a meta-analysis on SNHGs and HCC prognosis, but there were only five SNHGs included. In recent years, studies involving more SNHGs about HCC have emerged. So for further knowledge, we carried out a meta-analysis on the prognostic value and clinicopathological features of the SNHG family in HCC.

## 2. Methods

### 2.1. Literature searching strategies

We prospectively registered the meta-analysis on PROSPERO (CRD42022370591). To filter all pertinent studies up to October 2022, Embase, Web of Science, PubMed, and Cochrane Library were searched online. The following keywords were used in the search: ((Small nucleolar RNA host gene) OR SNHG OR (LncRNA SNHG)) AND ((Hepatocellular Carcinoma) OR (Hepatoma) OR (Liver Cell Carcinoma) OR (Liver Cancer)) AND (prognostic OR prognosis OR outcome). To make sure that all appropriate studies were included, we also manually searched pertinent original article references. Two researchers independently decided which publications should be included and excluded.

### 2.2. Inclusion and exclusion criteria

Inclusion criteria: (A) patients were definitely diagnosed as HCC, and their SNHG expression was detected by specific methods, (B) “High SNHG” and “low SNHG” groups were created. (C) the correlation between SNHG and prognosis along with clinical features were reported, (D) there was enough data in the studies to generate HRs and 95% CIs.

Exclusion criteria: (A) replicated studies, (B) review, meta, raw letter, cellular, and animal studies, (C) irrelevant topics, (D) inadequate information for calculation of HRs and 95% CIs.

### 2.3. Data Extraction and Quality Assessment

Data were separately extracted by two researchers using inclusion and exclusion criteria, and any differences were solved by discussion. We extracted the following information: the first author, study country, publication year, sample size, sample type, SNHG type, method of SNHG detection, the cut-off value of SNHG expression, endpoints, extract method of HR, HRs and 95% CIs of OS/RFS/DFS reported or estimated from survival curves, and clinical characteristics including age, sex, lymphatic metastasis, vein invasion, tumor size, tumor stage, histologic grade. If only Kaplan-Meier curves were given, we could calculate HR and 95% CI with the Engage Digitizer v11.1 software based on the strategy suggested by Tierney [13].

The Newcastle-Ottawa Quality Assessment Scale (NOS) was employed to examine the quality of the included studies. The NOS scale [14] considers inclusion, outcome, and comparability, with scores from 0 to 9. Studies with a score of 6 or higher were included in the meta-analysis.

### 2.4. Statistical Analysis

The STATA 15.0 software was used to conduct this meta-analysis of all eligible studies. The relationship between SNHG expression and the prognosis of HCC patients was evaluated by hazard ratios (HRs) and 95% confidence intervals (CIs). Additionally, odds ratios (ORs) and 95% CIs were used to analyze the association between clinicopathological traits and SNHG. We analyzed heterogeneity between studies with the Chi-square Q test and I^2^ statistic. The random-effects model was applied if there was significant heterogeneity (χ^2^ test P < 0.1 or I^2^ > 50%); otherwise, a fixed-effects model was utilized. A sensitivity analysis was conducted to check whether the findings were stable. Begg’s and Egger’s tests were used to assess publication bias. Significant differences were defined as P-values < 0.05.

## 3. Results

### 3.1. Search Process and Features of Included Literature

The detailed search and inclusion process is shown in Fig 1. There were 429 pieces of literature found overall in the initial search. We chose 46 studies strictly based on the criteria for inclusion and exclusion. Then, we further excluded 21 studies upon reviewing all full text of the remaining 46 articles. Ultimately, this meta-analysis included 25 publications (2314 patients) [15-39]. The sample size was between 40 and 160, with publication years ranging from 2016 to 2022. Except for one [37], all of the listed studies were carried out in China. These eligible studies included 2314 patients with 13 types of SNHGs, involving SNHG1 [18,34], SNHG3 [19,28], SNHG6 [15,20], SNHG7 [26,27,30,31], SNHG9 [37], SNHG10 [24], SNHG11 [29], SNHG12 [21], SNHG15 [17,33], SNHG16 [23,25,32,39], SNHG17 [36], SNHG18 [38], SNHG20 [16,22], and SNHG22 [35]. Based on SNHG expression, the enrolled patients were divided into the “high SNHG” or “low SNHG” group. Apart from one study that identified SNHG expression in venous blood, all of the research detected SNHG expression in tissue by qRT-PCR or ISH [37]. The correlation between SNHG and OS was explored in every study. Of them, eight studies also reported the correlation between SNHG and DFS/RFS. 13 studies gave HR values directly, while Kaplan-Meier curves were available for calculating HR values in the remaining studies. All included studies had NOS scores ≥ 6. The characteristics of the included literature are detailed in Table 1.

**Table 1.**
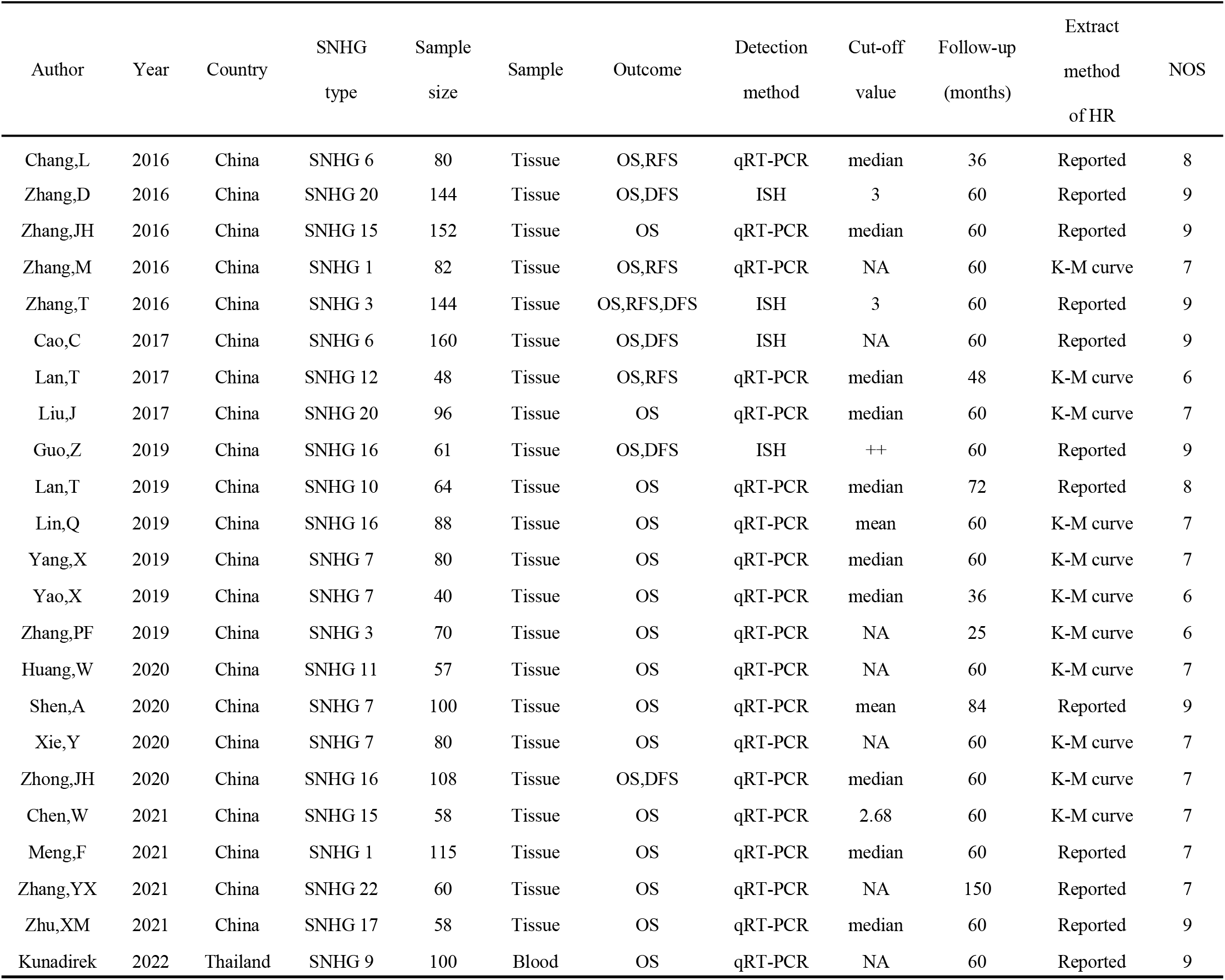
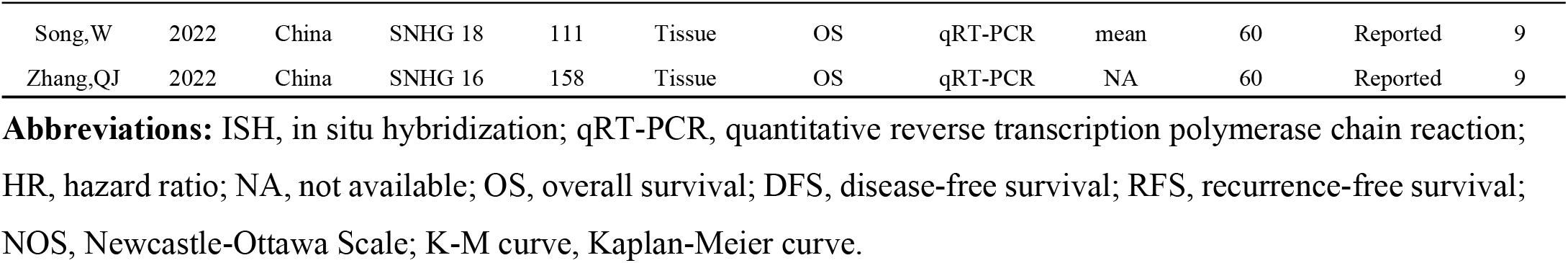
Details of included studies.

**Figure 1.**
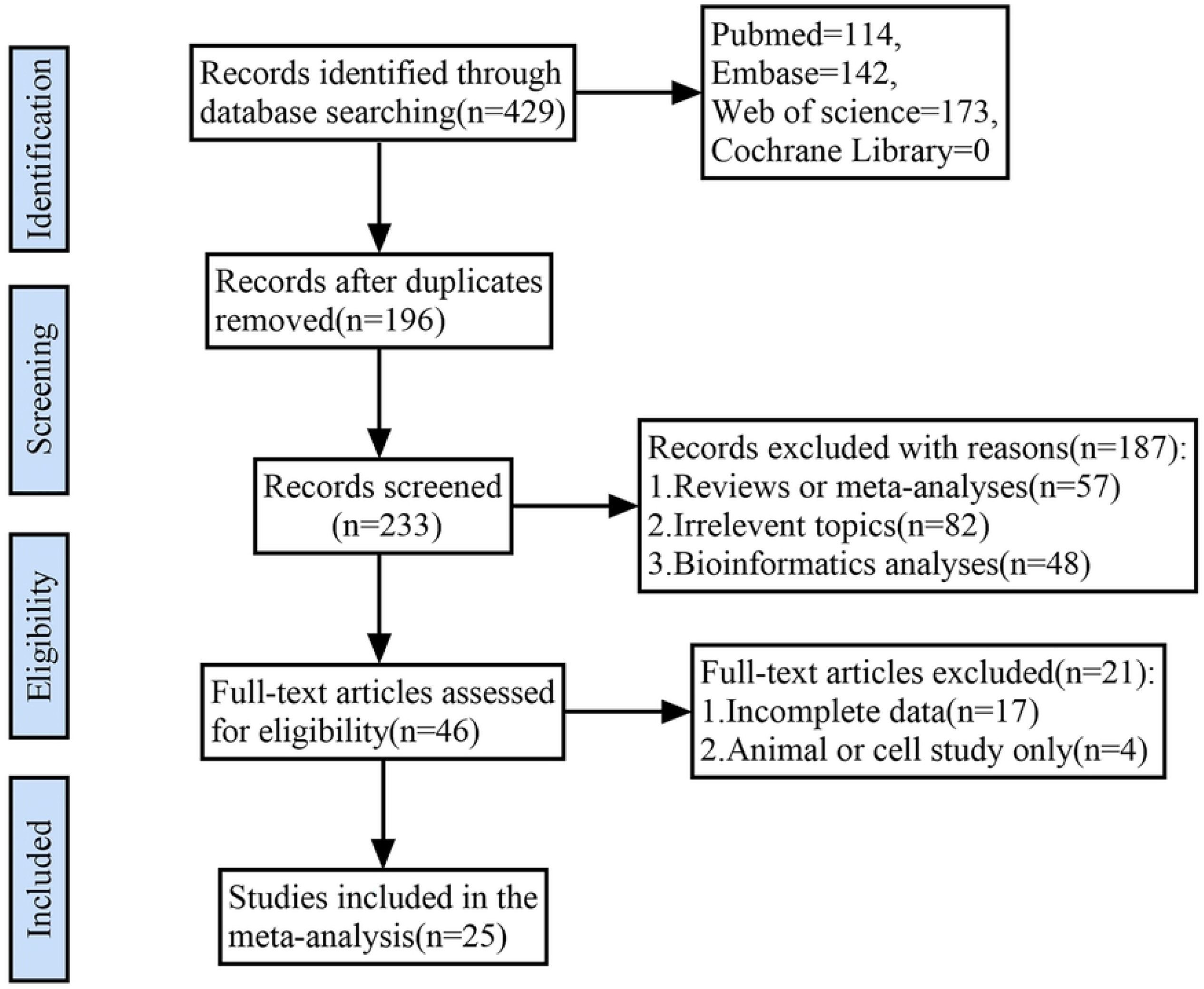
PRISMA flowchart of the selection process.

### 3.2. The Correlation on SNHG Expression and HCC Prognosis

#### 3.2.1. SNHG Expression and Overall Survival (OS)

To analyze the relationship of SNHG expression with OS, we selected 25 relevant studies with 2314 participants. The combined results showed a significant association between higher SNHG expression and shorter OS (HR: 2.22, 95% CI: 1.77-2.78, P: 0.000, S1 Fig). The HR and 95% CI of OS were analyzed using a random-effects model due to the heterogeneity that existed among the studies (I^2^: 70.5%, P: 0.000). Sensitivity analysis shown in S2 Fig revealed that “Lan T 2019” greatly influenced the stability of the results, which may be a possible source of heterogeneity. Then, after removing “Lan T 2019”, the new results showed no heterogeneity (I^2^: 0.00%, P: 0.896), indicating that Lan T 2019 was the source of heterogeneity (Fig 2a). As a result, we finally included 24 qualified studies with 2250 patients. A fixed-effects model determined that the total HR and 95% CI were significant statistically and showed that patients with up-regulated SNHGs had a higher risk of having a short OS (HR: 2.33, 95% CI: 2.02-2.69; P: 0.000). The results appeared to be trustable, as demonstrated in the sensitivity analysis of Fig 2. Furthermore, in order to carry out a subgroup analysis of OS, all included patients were grouped according to extract method of HR, sample size, SNHG type, and follow-up time (Table 2). Increased expression of SNHG1, 3, 6, 7, 15, 16, and 20 was substantially connected with poor OS in the SNHG type subgroup, as shown in S3 Fig, and other SNHGs were as well. We came to the conclusion that increased expression of SNHGs could result in worse OS both in the group of reported and survival curve when the studies were categorized using the HR extraction approach (reported: HR: 2.44, 95% CI: 2.05-2.90, P: 0.000; survival curve: HR: 2.12, 95% CI: 1.64-2.73, P: 0.000). In addition, we discovered that HCC patients with a worse prognosis had higher levels of SNHG expression regarding follow-up time and sample size. The mentioned data supported that SNHGs in HCC patients could serve as prognostic markers for the intervention of OS.

**Table 2.**
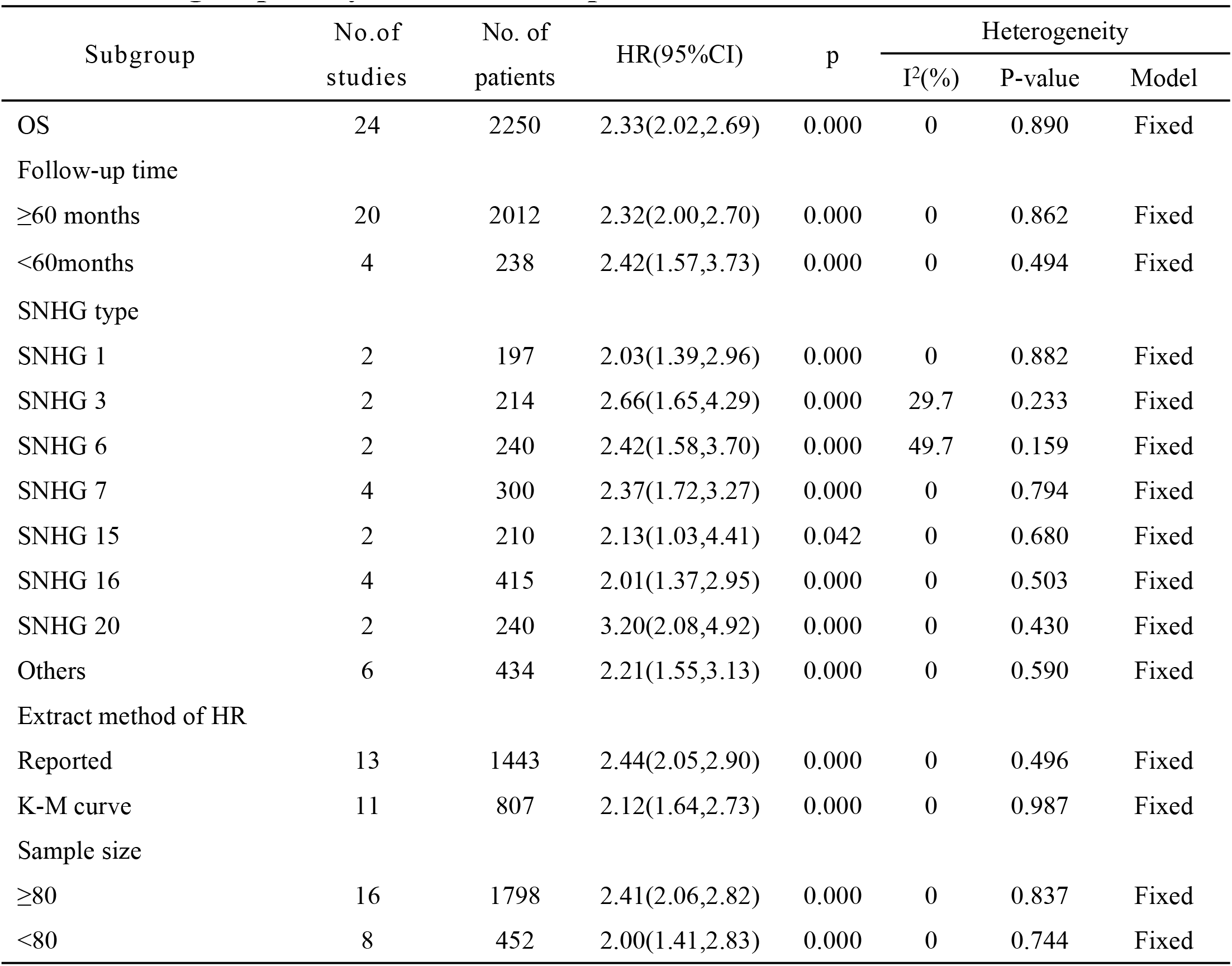
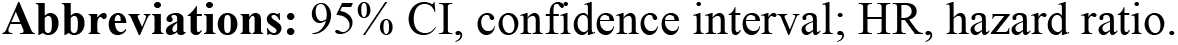
Subgroup analysis of SNHG expression for OS.

| Subgroup | No. of studies | No. of patients | HR(95%CI) | p | Heterogeneity |  |  |
| --- | --- | --- | --- | --- | --- | --- | --- |
|  |  |  |  |  | I<sup>2</sup>(%) | P-value | Model |
| OS | 24 | 2250 | 2.33(2.02,2.69) | 0.000 | 0 | 0.890 | Fixed |
| Follow-up time |  |  |  |  |  |  |  |
| ≥60 months | 20 | 2012 | 2.32(2.00,2.70) | 0.000 | 0 | 0.862 | Fixed |
| <60months | 4 | 238 | 2.42(1.57,3.73) | 0.000 | 0 | 0.494 | Fixed |
| SNHG type |  |  |  |  |  |  |  |
| SNHG 1 | 2 | 197 | 2.03(1.39,2.96) | 0.000 | 0 | 0.882 | Fixed |
| SNHG 3 | 2 | 214 | 2.66(1.65,4.29) | 0.000 | 29.7 | 0.233 | Fixed |
| SNHG 6 | 2 | 240 | 2.42(1.58,3.70) | 0.000 | 49.7 | 0.159 | Fixed |
| SNHG 7 | 4 | 300 | 2.37(1.72,3.27) | 0.000 | 0 | 0.794 | Fixed |
| SNHG 15 | 2 | 210 | 2.13(1.03,4.41) | 0.042 | 0 | 0.680 | Fixed |
| SNHG 16 | 4 | 415 | 2.01(1.37,2.95) | 0.000 | 0 | 0.503 | Fixed |
| SNHG 20 | 2 | 240 | 3.20(2.08,4.92) | 0.000 | 0 | 0.430 | Fixed |
| Others | 6 | 434 | 2.21(1.55,3.13) | 0.000 | 0 | 0.590 | Fixed |
| Extract method of HR |  |  |  |  |  |  |  |
| Reported | 13 | 1443 | 2.44(2.05,2.90) | 0.000 | 0 | 0.496 | Fixed |
| K-M curve | 11 | 807 | 2.12(1.64,2.73) | 0.000 | 0 | 0.987 | Fixed |
| Sample size |  |  |  |  |  |  |  |
| ≥80 | 16 | 1798 | 2.41(2.06,2.82) | 0.000 | 0 | 0.837 | Fixed |
| <80 | 8 | 452 | 2.00(1.41,2.83) | 0.000 | 0 | 0.744 | Fixed |

**Abbreviations:** 95% CI, confidence interval; HR, hazard ratio.

**Figure 2.**
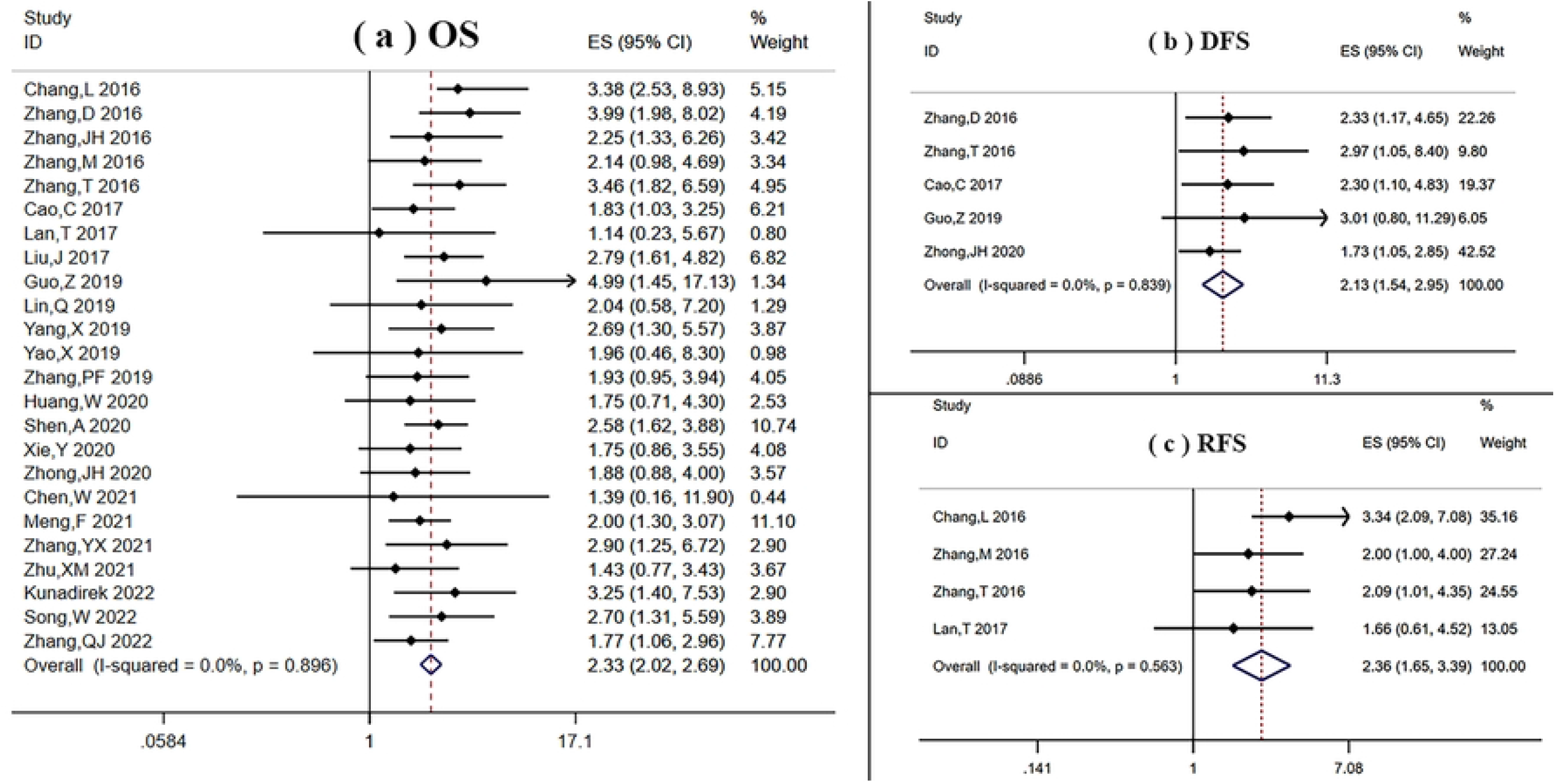
Forest plots of the relationship on SNHG expression and prognosis after deleting “Lan T 2019”: (a) OS; (b) DFS; (c) RFS.

#### 3.2.2. SNHG Expression and Recurrence Free Survival (RFS) or Disease Free Survival (DFS)

Figs 2b and 2c depict the relationship between SNHG expression and RFS, DFS respectively. Four studies [15,18,19,21] consisting of 354 patients demonstrated the utility of SNHGs as predictive biomarkers for RFS in HCC (HR: 2.36, 95% CI: 1.65-3.39, P: 0.000). Meanwhile, elevated expression of SNHG in HCC was likewise significantly related to poor DFS, according to five studies by Zhang et al. [16,19,20,23,31] (HR: 2.13, 95% CI: 1.54-2.95; P: 0.000).

### 3.3. The SNHG expression and clinicopathological characteristics

In the included papers, we conducted a meta-analysis to determine whether there was a correlation between the clinicopathological characteristics of HCC and SNHG expression. Table 3 presents the outcomes.

**Table 3.**
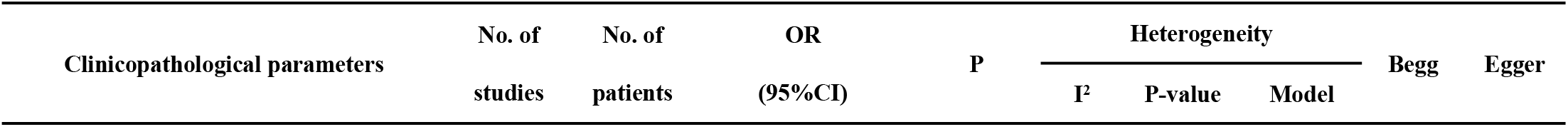
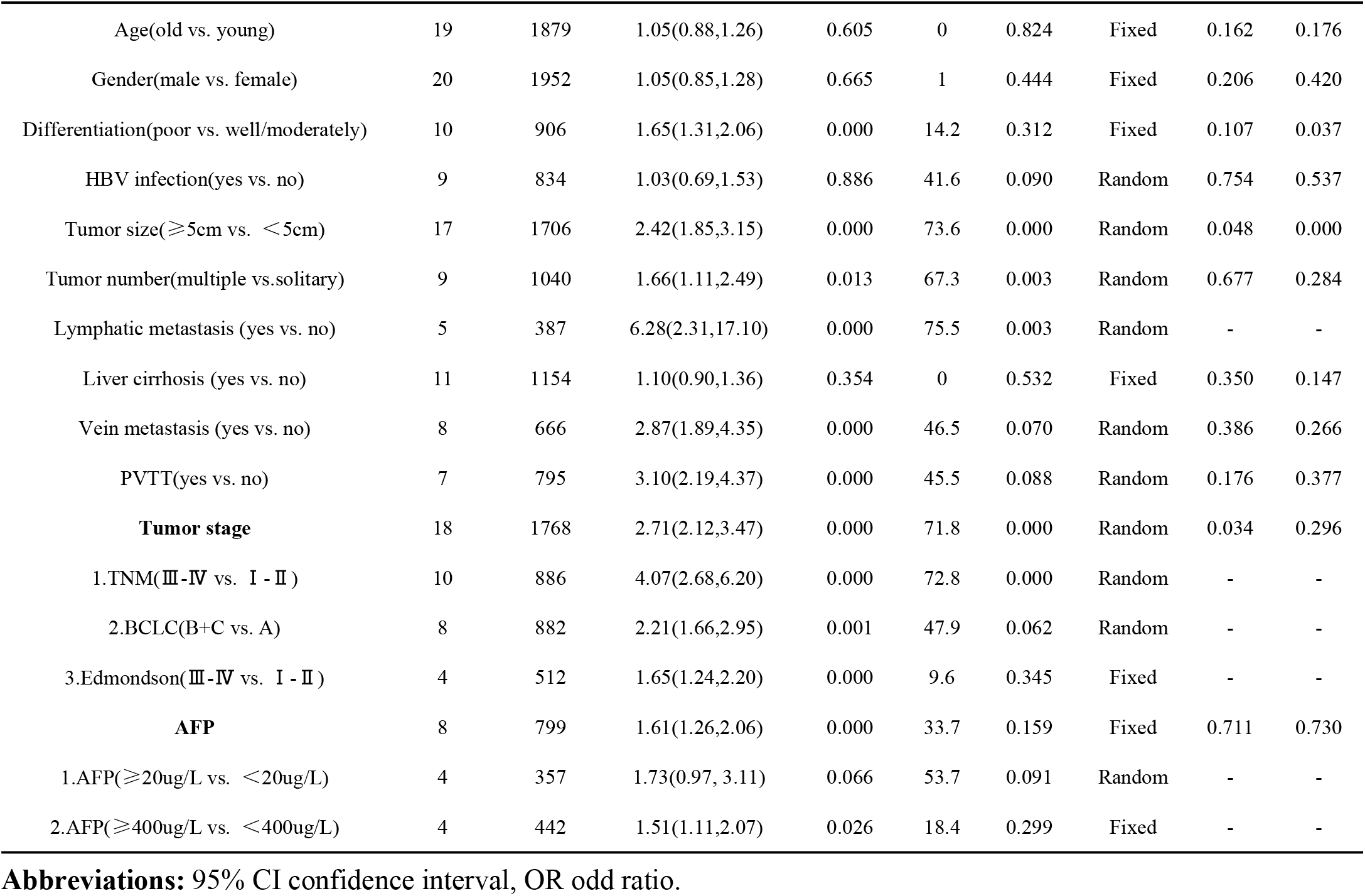
Meta-analysis results on the relationship between over-expressed SNHG and clinicopathological factors.

#### 3.3.1. SNHG and tumor size, number, and differentiation

Larger tumor size was found in the group of “high SNHG” in 17 studies (tumor diameter ≥5 cm vs. <5 cm, OR: 2.42, 95% CI: 1.85-3.15, P: 0.000; Figure 4a). Given the significant degree of heterogeneity (I^2^: 73.6%, P: 0.000), a random-effects model was used. To examine the association between SNHG and tumor number, 9 studies encompassing 1040 patients were analyzed with a random-effects model (I^2^: 67.3%, P: 0.003) (Fig 4b). The findings indicated that patients with multiple tumor numbers were probably accompanied by elevated SNHG levels (OR: 1.66; 95% CI: 1.11-2.49; P: 0.013). Meanwhile, adopting a fixed-effects model with no clear heterogeneity observed (I^2^: 14.2%, P: 0.312), 10 studies including 906 patients showed that HCC patients with SNHG overexpression had a higher likelihood of developing a low histological grade (Fig 4c OR: 1.65, 95% CI: 1.31-2.06; P: 0.000).

**Figure 3.**
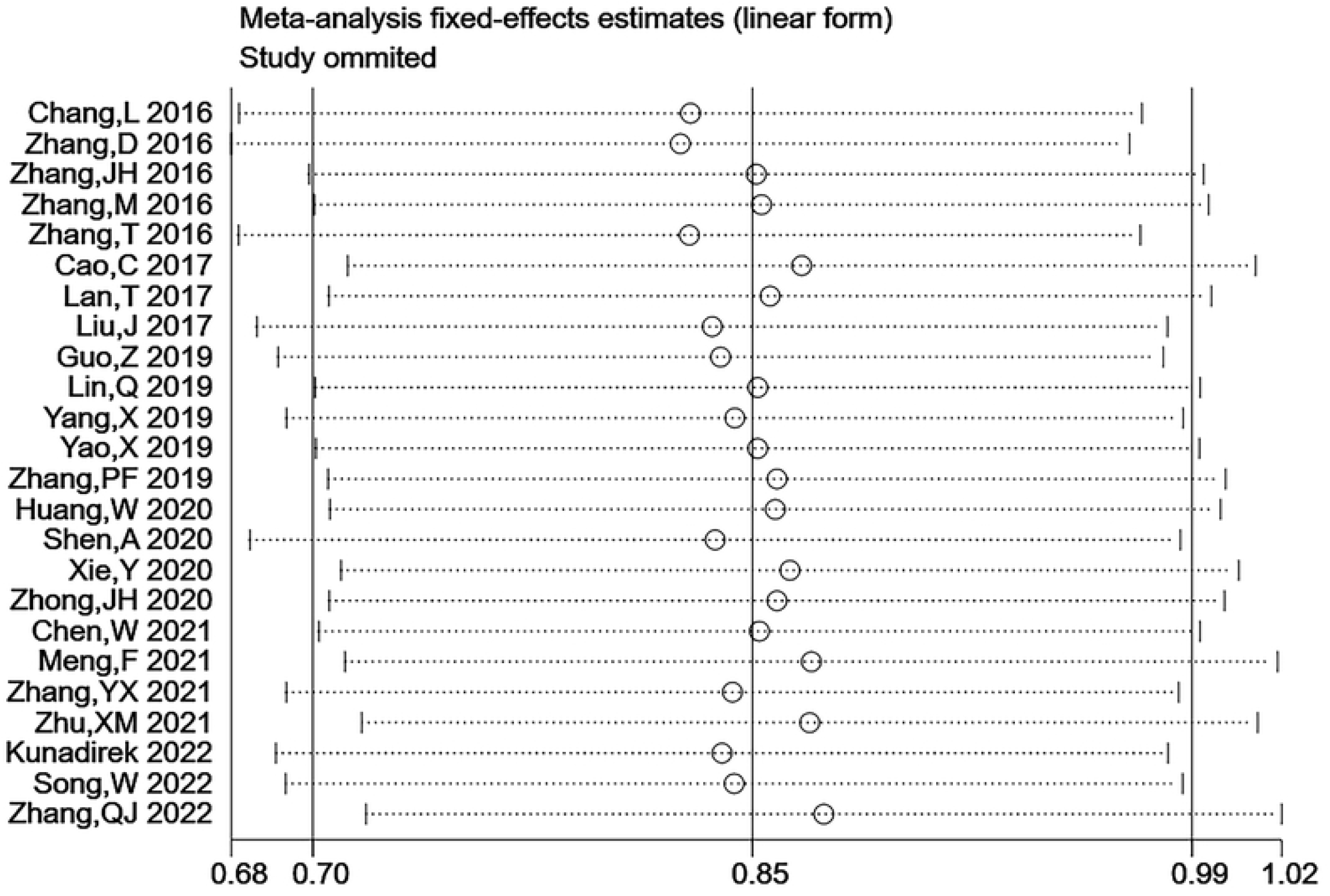
Sensitivity analysis of meta-analysis on SNHG and OS after deleting “Lan T 2019”.

**Figure 4.**
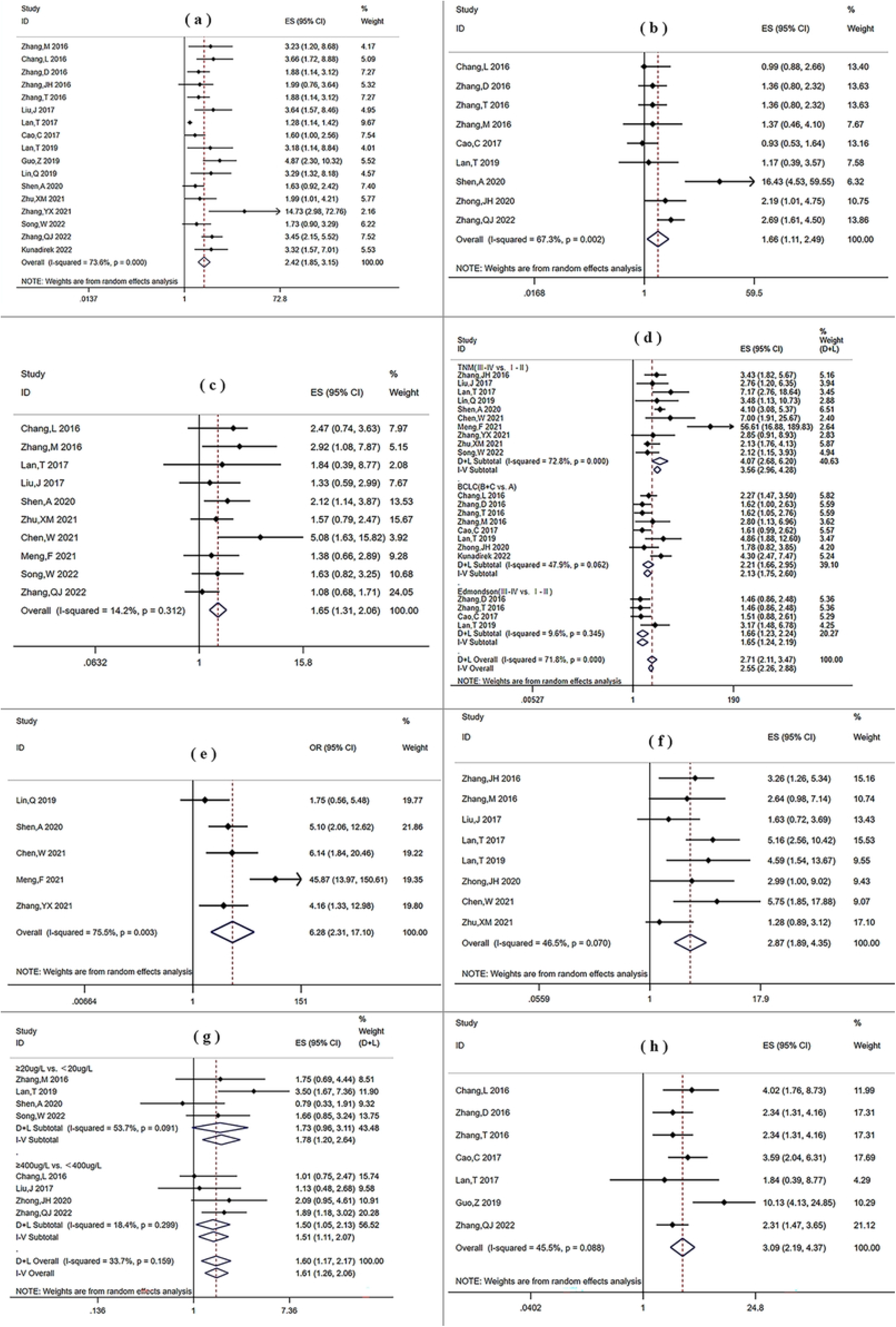
Forest plots on the association between SNHG expression and clinicopathological feactures: (a) tumor size; (b) tumor number; (c) differentiation; (d) tumor stage; (e) lymphatic metastasis; (f) vein Invasion; (g) AFP; (f) portal vein tumor thrombosis (PVTT).

#### 3.3.2. SNHG and Tumor Stage

We pooled the OR and 95% CI values from 18 research to analyze the relationship on SNHG expression and the HCC stage. A random-effects model was applied since the heterogeneity between studies was > 50% and the p-value < 0.1 (I^2^: 71.8%, P: 0.000). The combined OR and 95% CI had statistical significance (OR: 2.71, 95% CI: 2.12-3.47; P: 0.000), suggesting over-expressed SNHG was related with the advanced clinical stage(Fig 4d). When subgroup analysis was performed using stage criteria such as TNM, BCLC, and Edmondson stage, the same conclusion was reached (TNM(III-IV vs. I-II): OR: 4.07, 95% CI: 2.68-6.20; P: 0.000; BCLC(B+C vs. A): OR: 2.21, 95%CI: 1.66-2.95; P: 0.001; Edmondson(III-IV vs. I-II): OR: 1.65, 95% CI: 1.24-2.20; P: 0.000).Thus, overexpression of SNHG was likely to raise the risk of advanced-stage HCC.

#### 3.3.3. SNHG and Lymphatic Metastasis and Vein Invasion

There were 5 studies involving 387 participants examined the association of SNHG expression and lymphatic metastasis. The total OR and 95% CI were calculated using a random-effects model due to the heterogeneity among studies (I^2^: 75.5%, P: 0.003; Fig 4e). According to the findings, early lymph node metastasis was linked to upregulated SNHG expression in HCC patients (OR: 6.28, 95% CI: 2.31-17.10; P: 0.000). Meanwhile, patients with high SNHG expression had a higher risk of vein invasion than those with low expression, according to the pooled results from 8 high-quality articles (Fig 4f, OR: 2.87, 95% CI: 1.89-4.35, P: 0.000) by a random-effects model (I^2^: 46.5%, P: 0.070).

#### 3.3.4. SNHG with Portal Vein Tumor Thrombosis (PVTT) and AFP

The relationship on SNHG expression and the AFP value of patients was explored in 8 studies. No notable heterogeneity among the research was found, thus the fixed-effects model with an OR of 1.61 (95% CI: 1.26-2.06, P: 0.000) (Fig 4g), showed that increased levels of SNHGs tended to be accompanied by higher AFP value in HCC. Furthermore, 4 studies of them had an AFP cut-off value of 20ug/L and the others were 400ug/L. AFP >400ug/L was substantially related to high expression of SNHG, while AFP >20ug/L exhibited no noticeable difference, according to the subgroup analysis carried out depending on the cut-off values (AFP(≥20ug/L vs. 20ug/L): OR: 1.73, 95% CI: 0.97-3.11, P: 0.066; AFP(≥400ug/L vs. <400ug/L): OR: 1.51, 95% CI: 1.11-2.07, P: 0.026). Then, to see if SNHG expression was related to combined portal vein tumor thrombosis (PVTT) in HCC patients, we analyzed 7 studies involving 795 patients(Fig 4h). The random-effects model we chose revealed that PVTT was associated with increased SNHG expression in HCC patients due to the high heterogeneity among studies(I^2^: 45.5%, P: 0.088; OR: 3.10, 95% CI: 2.19-4.37, P: 0.000). Besides, the relationship between SNHG expression and other clinical characteristics was also explored such as age (OR: 1.05, 95% CI: 0.88-1.26, P: 0.605), gender (OR: 1.05, 95% CI: 0.85-1.28, P: 0.665), HBV infection (OR: 1.03, 95% CI: 0.69-1.53, P: 0.886) and cirrhosis (OR: 1.10, 95% CI: 0.90-1.36, P: 0.354). However, no significant associations were found.

### 3.3. Publication Bias

The funnel plot, Begg’s test, and Egger’s test were used to examine potential publication bias. The meta-analysis for OS did not exhibit any publication bias, according to Begg’s test and Egger’s test (P: 0.862 and P: 0.850, respectively), and Begg’s funnel plot in Fig 5 was generally symmetrical. In addition, we assessed publication bias in terms of clinicopathological characteristics, including age, gender, histological grade, vein invasion, cirrhosis, tumor number, tumor size, HBV infection, PVTT, tumor stage, and AFP (Table III).

**Fig 5.**
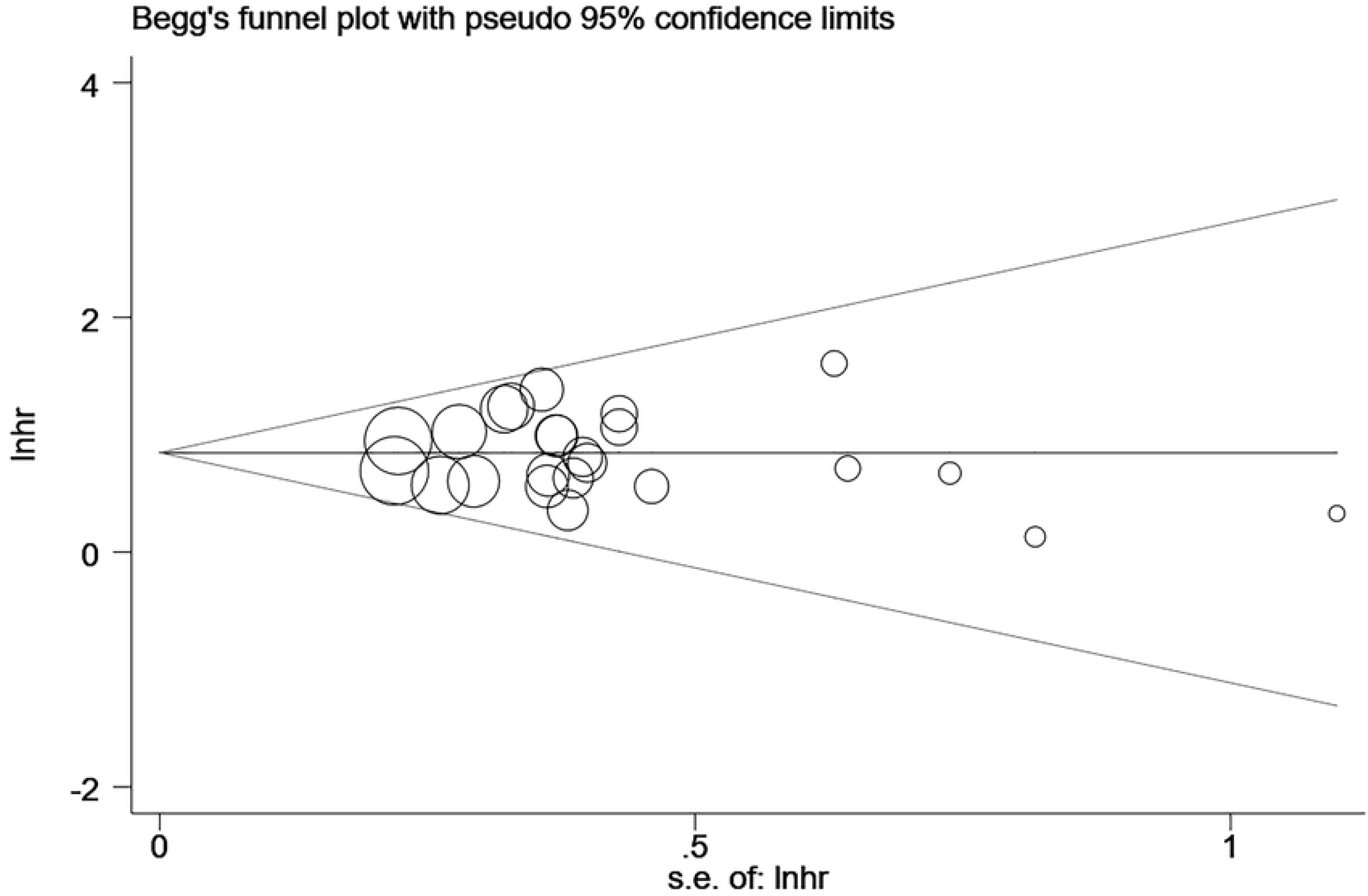
Begg’s funnel plot of publication bias for OS. Each circle represents an independent study.

## 4. Discussion

LncRNAs are RNA molecules with a length more than 200 nucleotides that are incapable of being translated into proteins. Recent research has shown that lncRNAs regulate gene expression, which have a profound impact on various physiological and pathological processes. In particular, they can function as oncogenes or anti-oncogenes that directly or indirectly control signaling pathways associated with tumors and influence tumor progression [40]. Due to the elevated activation of carcinogenic lncRNAs in various cancers in adjacent tissues, concern over lncRNAs as diagnostic or prognostic markers is growing [41]. Similarly, The SNHGs may also be promising indicators for the detection and prognosis of cancer according to many research.

It has been extensively studied how SNHGs contribute to the growth of HCC. SNHGs can influence the biologically malignant behavior of tumors through acting as endogenous RNAs (ceRNAs) to adsorb numerous miRNAs, directly binding to and upregulating mRNA, interacting with transcription factors to activate transcription, and activating the signaling pathways such as Wnt/β-catenin that are primarily part in tumor growth. They significantly affected the outcome of HCC patients by promoting growth and metastasis or interfering with apoptosis and autophagy in HCC [11]. Zhang et al [18] stated SNHG1 drastically improved the proliferation and migration of HCC by suppressing the activity of p53 and its target genes, which also inhibited the apoptosis process. According to the study by Xie et al [31], SNHG7, which was highly expressed in HCC, might compete with miR-9-5p as a ceRNA. As a result, it could upregulate the activity of the CNNM1, which encouraged the growth of HCC in turn. The majority of SNHGs functioned as ceRNAs to promote malignancy in HCC. In addition, SNHGs also played important roles in HCC via additional mechanisms. For example, in HCC, SNHG3 [28], SNHG7 [27], and SNHG20 [22] promoted epithelial-to-mesenchymal transition (EMT), and SNHG16 [39] significantly activated the ECM-receptor interaction pathway, and so on. To sum up, mechanisms of SNHGs varied in HCC.

Therefore, we carried out a further analysis to determine if SNHGs were valuable in the prognosis of HCC.

In our analysis to summarize the relationship on prognosis and clinicopathological parameters with SNHG in HCC, a total of 25 eligible papers satisfying inclusion criteria were taken into account. Patients with high SNHG expression had a higher chance of undergoing shorter OS, DFS, and RFS compared to those with low expression. Additionally, the association between SNHG and OS under various situations was further investigated with subgroup analysis. All pooled results showed that SNHG overexpression was related to worse OS of HCC in subgroups of SNHG type, extract method, follow-up time, and sample size.

The correlation on SNHG expression and various clinicopathological features in HCC was also explored at the same time. Patients who had high levels of SNHGs expression were inclined to have larger tumor size, multiple tumors, worse histologic grades, more advanced tumor stage, positive lymphatic metastasis, vein invasion, PVTT, and AFP values > 400ug/L. In contrast, SNHG expression had no bearing on a range of other clinical characteristics, including age, gender, HBV infection, and cirrhosis. Taking into account all of these findings, lncRNA SNHGs played a part in the emergence of HCC and had the potential to grow into a biomarker for the prognosis. Our findings were consistent with a meta-analysis [12] on SNHG and HCC prognosis in 2017, which covered only 5 papers at that time. In recent years, more studies on this subject have been published. Then, we reanalyzed the association between SNHG and clinical features along with the prognosis of HCC, which appeared more convincing and detailed.

However, we should acknowledge some limitations of the study. Firstly, the results couldn’t be generalized to other nations because all but one of the research included in it were from China, which could have caused publication bias. Secondly, there was a chance of errors because the HRs and 95% CIs were partially extracted indirectly from the survival curves. Thirdly, the cut-off values of SNHG varied between research, which could have an impact on the relevant outcomes. Lastly, only a portion of SNHGs was involved in our analysis due to insufficient research and sample size.

## 5. Conclusion

To summarize, we concluded from this meta-analysis that upregulation of SNHG related to malignant clinicopathological features of HCC and had potential as a prognostic biomarker. Future large-scale, multicenter studies will be necessary to confirm our findings.

## Supporting information

**S1 Fig. Forest plot for the relationship between SNHG expression and OS before deleting “Lan T 2019”**.

**S2 Fig. Sensitivity analysis for meta-analysis of SNHG and OS before deleting “Lan T 2019”**.

**S3 Fig. Forest plots of hazard ratios (HRs) for the relationship between SNHG**

**expression and OS:** (a) SNHG type; (b)extract method of HR; (c) follow-up time; (d) sample size.

**S1 Table. NOS quality scores of included studies**.

## Acknowledgements

NA.

## Data availability

All relevant data are within the manuscript and its Supporting Information files.

## Author contributions

**Conceptualization:** Lihua Yang

**Data curation:** Zhiwen Zhao

**Resources:** Meng Huang

**Supervision:** Lihua Yang

**Writing - original draft:** Meng Huang

**Writing - review & editing:** Lihua Yang

## References

[1] Sung H, Ferlay J, Siegel RL, Laversanne M, Soerjomataram I, Jemal A, et al. Global Cancer Statistics 2020: GLOBOCAN Estimates of Incidence and Mortality Worldwide for 36 Cancers in 185 Countries. CA Cancer J Clin. 2021; 71: 209–249.

[2] McGlynn KA, Petrick JL, El-Serag HB. Epidemiology of Hepatocellular Carcinoma. Hepatology. 2021; 73 Suppl 1: 4–13.

[3] Chen Z, Xie H, Hu M, Huang T, Hu Y, Sang N, et al. Recent progress in treatment of hepatocellular carcinoma. Am J Cancer Res. 2020; 10: 2993–3036.

[4] Xiong Y, Cao P, Lei X, Tang W, Ding C, Qi S, et al. Accurate prediction of microvascular invasion occurrence and effective prognostic estimation for patients with hepatocellular carcinoma after radical surgical treatment. World J Surg Oncol. 2022; 20: 328.

[5] Yao ZT, Yang YM, Sun MM, He Y, Liao L, Chen KS, et al. New insights into the interplay between long non-coding RNAs and RNA-binding proteins in cancer. Cancer Commun (Lond). 2022; 42: 117–140.

[6] Lin YH. Crosstalk of lncRNA and Cellular Metabolism and Their Regulatory Mechanism in Cancer. Int J Mol Sci. 2020; 21: 2947.

[7] Huang Z, Zhou JK, Peng Y, He W, Huang C. The role of long noncoding RNAs in hepatocellular carcinoma. Mol Cancer. 2020; 19: 77.

[8] Zimta AA, Tigu AB, Braicu C, Stefan C, Ionescu C, Berindan-Neagoe I. An Emerging Class of Long Non-coding RNA With Oncogenic Role Arises From the snoRNA Host Genes. Front Oncol. 2020; 10: 389.

[9] Biagioni A, Tavakol S, Ahmadirad N, Zahmatkeshan M, Magnelli L, Mandegary A, Samareh Fekri H, Asadi MH, Mohammadinejad R, Ahn KS. Small nucleolar RNA host genes promoting epithelial-mesenchymal transition lead cancer progression and metastasis. IUBMB Life. 2021; 73: 825–842.

[10] Saeinasab M, Atlasi Y, M Matin M. Functional role of lncRNAs in gastrointestinal malignancies: the peculiar case of small nucleolar RNA host gene family. FEBS J. 2022; article 16668.

[11] Li Y, Wang X, Chen S, Wu B, He Y, Du X, Yang X. Long non-coding RNA small nucleolar RNA host genes: functions and mechanisms in hepatocellular carcinoma. Mol Biol Rep. 2022; 49: 2455–2464.

[12] Li J, Gao J, Kan A, Hao T, Huang L. SNHG and UCA1 as prognostic molecular biomarkers in hepatocellular carcinoma: recent research and meta-analysis. Minerva Med. 2017; 108: 568–574.

[13] Tierney JF, Stewart LA, Ghersi D, Burdett S, Sydes MR. Practical methods for incorporating summary time-to-event data into meta-analysis. Trials. 2007; 8: 16.

[14] Stang A. Critical evaluation of the Newcastle-Ottawa scale for the assessment of the quality of nonrandomized studies in meta-analyses. Eur J Epidemiol. 2010; 25: 603–5.

[15] Chang L, Yuan Y, Li C, Guo T, Qi H, Xiao Y, Dong X, Liu Z, Liu Q. Upregulation of SNHG6 regulates ZEB1 expression by competitively binding miR-101-3p and interacting with UPF1 in hepatocellular carcinoma. Cancer Lett. 2016; 383: 183–194.

[16] Zhang D, Cao C, Liu L, Wu D. Up-regulation of LncRNA SNHG20 Predicts Poor Prognosis in Hepatocellular Carcinoma. J Cancer. 2016; 7: 608–17.

[17] Zhang JH, Wei HW, Yang HG. Long noncoding RNA SNHG15, a potential prognostic biomarker for hepatocellular carcinoma. Eur Rev Med Pharmacol Sci. 2016; 20: 1720–4.

[18] Zhang M, Wang W, Li T, Yu X, Zhu Y, Ding F, Li D, Yang T. Long noncoding RNA SNHG1 predicts a poor prognosis and promotes hepatocellular carcinoma tumorigenesis. Biomed Pharmacother. 2016; 80: 73–79.

[19] Zhang T, Cao C, Wu D, Liu L. SNHG3 correlates with malignant status and poor prognosis in hepatocellular carcinoma. Tumour Biol. 2016; 37: 2379–85.

[20] Cao C, Zhang T, Zhang D, Xie L, Zou X, Lei L, Wu D, Liu L. The long non-coding RNA, SNHG6-003, functions as a competing endogenous RNA to promote the progression of hepatocellular carcinoma. Oncogene. 2017; 36: 1112–1122.

[21] Lan T, Ma W, Hong Z, Wu L, Chen X, Yuan Y. Long non-coding RNA small nucleolar RNA host gene 12 (SNHG12) promotes tumorigenesis and metastasis by targeting miR-199a/b-5p in hepatocellular carcinoma. J Exp Clin Cancer Res. 2017; 36: 11.

[22] Liu J, Lu C, Xiao M, Jiang F, Qu L, Ni R. Long non-coding RNA SNHG20 predicts a poor prognosis for HCC and promotes cell invasion by regulating the epithelial-to-mesenchymal transition. Biomed Pharmacother. 2017; 89: 857–863.

[23] Guo Z, Zhang J, Fan L, Liu J, Yu H, Li X, Sun G. Long Noncoding RNA (lncRNA) Small Nucleolar RNA Host Gene 16 (SNHG16) Predicts Poor Prognosis and Sorafenib Resistance in Hepatocellular Carcinoma. Med Sci Monit. 2019; 25: 2079–2086.

[24] Lan T, Yuan K, Yan X, Xu L, Liao H, Hao X, Wang J, Liu H, Chen X, Xie K, Li J, Liao M, Huang J, Zeng Y, Wu H. LncRNA SNHG10 Facilitates Hepatocarcinogenesis and Metastasis by Modulating Its Homolog SCARNA13 via a Positive Feedback Loop. Cancer Res. 2019; 79: 3220–3234.

[25] Lin Q, Zheng H, Xu J, Zhang F, Pan H. LncRNA SNHG16 aggravates tumorigenesis and development of hepatocellular carcinoma by sponging miR-4500 and targeting STAT3. J Cell Biochem. 2019; 120: 11604–11615.

[26] Yang X, Sun L, Wang L, Yao B, Mo H, Yang W. LncRNA SNHG7 accelerates the proliferation, migration and invasion of hepatocellular carcinoma cells via regulating miR-122-5p and RPL4. Biomed Pharmacother. 2019; 118: 109386.

[27] Yao X, Liu C, Liu C, Xi W, Sun S, Gao Z. lncRNA SNHG7 sponges miR-425 to promote proliferation, migration, and invasion of hepatic carcinoma cells via Wnt/β-catenin/EMT signalling pathway. Cell Biochem Funct. 2019; 37: 525–533.

[28] Zhang PF, Wang F, Wu J, Wu Y, Huang W, Liu D, Huang XY, Zhang XM, Ke AW. LncRNA SNHG3 induces EMT and sorafenib resistance by modulating the miR-128/CD151 pathway in hepatocellular carcinoma. J Cell Physiol. 2019; 234: 2788–2794.

[29] Huang W, Huang F, Lei Z, Luo H. LncRNA SNHG11 Promotes Proliferation, Migration, Apoptosis, and Autophagy by Regulating hsa-miR-184/AGO2 in HCC. Onco Targets Ther. 2020; 13: 413–421.

[30] Shen A, Ma J, Hu X, Cui X. High expression of lncRNA-SNHG7 is associated with poor prognosis in hepatocellular carcinoma. Oncol Lett. 2020; 19: 3959–3963.

[31] Xie Y, Wang Y, Gong R, Lin J, Li X, Ma J, Huo L. SNHG7 Facilitates Hepatocellular Carcinoma Occurrence by Sequestering miR-9-5p to Upregulate CNNM1 Expression. Cancer Biother Radiopharm. 2020; 35: 731–740.

[32] Zhong JH, Xiang X, Wang YY, Liu X, Qi LN, Luo CP, Wei WE, You XM, Ma L, Xiang BD, Li LQ. The lncRNA SNHG16 affects prognosis in hepatocellular carcinoma by regulating p62 expression. J Cell Physiol. 2020; 235: 1090–1102.

[33] Chen W, Huang L, Liang J, Ye Y, Yu S, Zhang Y. Long noncoding RNA small nucleolar RNA host gene 15 deteriorates liver cancer via microRNA-18b-5p/LIM-only 4 axis. IUBMB Life. 2021; 73: 349–361.

[34] Meng F, Liu J, Lu T, Zang L, Wang J, He Q, Zhou A. SNHG1 knockdown upregulates miR-376a and downregulates FOXK1/Snail axis to prevent tumor growth and metastasis in HCC. Mol Ther Oncolytics. 2021; 21: 264–277.

[35] Zhang Y, Lu C, Cui H. Long non-coding RNA SNHG22 facilitates hepatocellular carcinoma tumorigenesis and angiogenesis via DNA methylation of microRNA miR-16-5p. Bioengineered. 2021; 12: 7446–7458.

[36] Zhu XM, Li L, Ren LL, D. L, Wang YM. LncRNA SNHG17 predicts poor prognosis and promotes cell proliferation and migration in hepatocellular carcinoma. Eur Rev Med Pharmacol Sci. 2021; 25: 4219–4227.

[37] Kunadirek P, Pinjaroen N, Nookaew I, Tangkijvanich P, Chuaypen N. Transcriptomic Analyses Reveal Long Non-Coding RNA in Peripheral Blood Mononuclear Cells as a Novel Biomarker for Diagnosis and Prognosis of Hepatocellular Carcinoma. Int J Mol Sci. 2022; 23: 7882.

[38] Song W, Zhang X, Feng L, Lai Y, Li T, Zhang P. Downregulated lncRNA SNHG18 Suppresses the Progression of Hepatitis B Virus-Associated Hepatocellular Carcinoma and Meditates the Antitumor Effect of Oleanolic Acid. Cancer Manag Res. 2022; 14: 687–695.

[39] Zhang QJ, Li DZ, Lin BY, Geng L, Yang Z, Zheng SS. SNHG16 promotes hepatocellular carcinoma development via activating ECM receptor interaction pathway. Hepatobiliary Pancreat Dis Int. 2022; 21: 41–49.

[40] Zhang X, Hong R, Chen W, Xu M, Wang L. The role of long noncoding RNA in major human disease. Bioorg Chem. 2019; 92: 103214.

[41] Aprile M, Costa V, Cimmino A, Calin GA. Emerging role of oncogenic long noncoding RNA as cancer biomarkers. Int J Cancer. 2023; 152: 822–834.

